# Dynamics in Natural and Designed Elastins and Their Relation to Elastic Fiber Structure and Recoil

**DOI:** 10.1101/2020.07.14.202523

**Authors:** Ma. Faye Charmagne A. Carvajal, Jonathan M. Preston, Nour M. Jamhawi, T. Michael Sabo, Shibani Bhattacharya, James M. Aramini, Richard J. Wittebort, Ronald L. Koder

**Affiliations:** Department of Physics, The City College of New York, New York, New York 10031, United States; Department of Chemistry, University of Louisville, Louisville, Kentucky 40292, United States; Department of Medicine and the James Brown Cancer Center, University of Louisville School of Medicine, Louisville, Kentucky 40292, United States; Advanced Science Research Center, The City University of New York, New York, New York 10031, United States; Graduate Programs of Physics, Chemistry and Biochemistry, The Graduate Center of CUNY, New York, New York 10016, United States; The New York Structural Biology Center, New York, New York, 10031, United States

**Keywords:** IDP, minielastin, elastin, elastic recoil, dynamics, configurational entropy

## Abstract

Elastin fibers assemble in the extracellular matrix from the precursor protein tropoelastin and provide the flexibility and spontaneous recoil required for arterial function. Unlike many proteins, a structure-function mechanism for elastin has been elusive. We have performed detailed NMR relaxation studies of the dynamics of the minielastins **24x′** and **20x′** using solution NMR, and of purified bovine elastin fibers in the presence and absence of mechanical stress using solid state NMR. The low sequence complexity of the minielastins enables us to determine dynamical timescales and degrees of local ordering with residue-specific resolution in the cross-link and hydrophobic modules using NMR relaxation. We find an extremely high degree of disorder, with order parameters for the entirety of the hydrophobic domains near zero, resembling that of simple chemical polymers and less than the order parameters that have been observed in other intrinsically disordered proteins. We find that backbone order parameters in natural, purified elastin fibers are comparable to those found in **24x′** and **20x′** in solution. The difference in dynamics, compared to the minielastins, is that backbone correlation times are significantly slowed in purified elastin. Moreover, when elastin is mechanically stretched, the high chain disorder in purified elastin is retained - showing that any change in local ordering is below that detectable in our experiment. Combined with our previous finding of a 10-fold increase in the ordering of water when fully hydrated elastin fibers are stretched by 50%, these results support the hypothesis that stretch induced solvent ordering, i.e., the hydrophobic effect, is a key player in the elastic recoil of elastin as opposed to configurational entropy loss.

**SIGNIFICANCE:** Elastin is responsible for the spontaneous recoil of arterial walls that is necessary for cardiovascular function. Despite this critical role, the mechanism driving entropic recoil has remained unclear. Elastin is unusual in that it is intrinsically disordered in both soluble and fibrous forms. Using NMR, we have determined the timescales and amplitudes of dynamics in two soluble elastin mimetics and in relaxed and stretched states of purified bovine elastin fibers. Although dynamical timescales are different, both the soluble elastin mimetic and fibrillar elastin display an exceptionally high degree of disorder. No detectable increase in protein ordering was observed upon stretching, suggesting that entropic recoil is primarily driven by the hydrophobic effect and not configurational entropy loss.

## INTRODUCTION

Elastin, an extracellular matrix protein that is the principal elastic protein in vertebrates, is abundantly expressed in blood vessels, lung tissue, ligaments, and skin (1). The mature elastic matrix is formed when tropoelastin, one of the most hydrophobic proteins found in nature, is exported to the extracellular matrix and consecutively undergoes an oligomerization transition known as coacervation, followed by cross-linking via the enzymatic oxidation of lysyl ε-amino groups (2). The reversible entropic elasticity of fully matured elastin fibers in blood vessel walls is responsible for elastic energy storage during the cardiac cycle and the dampening of pulsatile flow in distal arteries via the Windkessel effect (3). Elastogenesis terminates in adolescence, and in the course of a human lifetime, arterial elastin undergoes in excess of 10^7^ stretching/contracting cycles. Oxidative damage accrued over human elastin’s lifetime reduces blood vessel compliance, resulting in hypertension, vascular calcification, ventricular hypertrophy, renal dysfunction and stroke (4). It is critical to understand the structural and dynamic origin of elastin’s entropic elasticity in order to understand how this oxidative structural damage leads to the pathogenesis and progression of these diseases.

Elastin proteins are organized in alternating proline-rich hydrophobic domains and alanine-rich cross-linking domains (5). The hydrophobic domains are quasi-repeats of three to seven amino acids rich in hydrophobic residues including proline, whereas the alanine-rich cross-linking domains are weakly helical (6) and present the cross-linking lysine residues in close proximity at *i* and *i+3* or *i+4* positions (7). The elastic function of elastin proteins primarily arises from the hydrophobic domains. These were for some time thought to have a stable repeating type II β-turn secondary structure (8). However, we have shown using nuclear magnetic resonance (NMR) analyses of both natural elastin fibers (9) and a series of simplified designed minielastin proteins (6) that these domains are intrinsically disordered in the unstressed relaxed state (9).

The entropically-driven recoil of a stretched disordered polymer can have two different origins: configurational entropy gain similar to that of vulcanized rubber (10), or the reduction of hydrophobic side chain exposure as the domain contracts and becomes more compact (11, 12). It is an important open question to what extent each of these mechanisms contribute to elastin function. In order to answer this question, we have performed detailed NMR studies of the dynamics of two designed minielastin proteins in solution - **24x′** and **20x′**- and of purified bovine elastin in the presence and absence of mechanical stress.

Like natural elastin, our designed minielastins have an alternating structure of hydrophobic modules, (APGVGV)_7_ or (VPGVGG)_5_ and cross-link modules, (DA_5_KA_2_KF) (see Scheme 1). Unlike natural elastin, **24x**′ and **20x**′ have identical repeats, which has allowed us to completely assign resonances in the proteins (6). In a protein with dynamic motion on these time scales, NMR chemical shifts reflect the time average chemical environment of each atom. In our earlier paper, we closely looked for chemical shift variation in the hydrophobic domains and found that chemical shifts for all but the outermost residues are identical and independent of whether the hydrophobic domain is in isolation, internal, or located at either protein terminus. This includes backbone (H_a_, C_a_, CO, HN and N) and side-chain atoms. Furthermore, varying the number of modules in the construct or changing the sequence of the cross-link modules does not change the chemical shifts. Given the narrow linewidths, we believe that the identical chemical shifts reflect identical ensemble behavior in every identical domain. Within the uncertainty of 2° chemical shifts, the observed 2° shifts are equivalent to random coil values in the hydrophobic modules and somewhat shifted toward α-helical values in the center of the cross-linker (6). All of these observations are consistent with a high degree of local dynamics. We also note that hydrophobic modules in natural elastin have an approximate repeat-like sequence. Combined with the reduced range of chemical shifts in IDP’s, complete residue specific NMR studies of dynamics in this type of protein are technically unfeasible and treating all repeats as dynamically equivalent is a reasonable and necessary approximation which has allowed us to determine both the dynamical timescale(s) *and the degree of local ordering* with residue-specific resolution in the cross-link and hydrophobic modules of **24x′** and **20x′** from solution NMR relaxation studies (R_1_, R_2_ and NOE) at three field strengths. We find an extremely high degree of disorder resembling that of some simple chemical polymers and comparable to only the smallest order parameters observed in intrinsically disordered natural proteins (13–15).

**Scheme 1.**
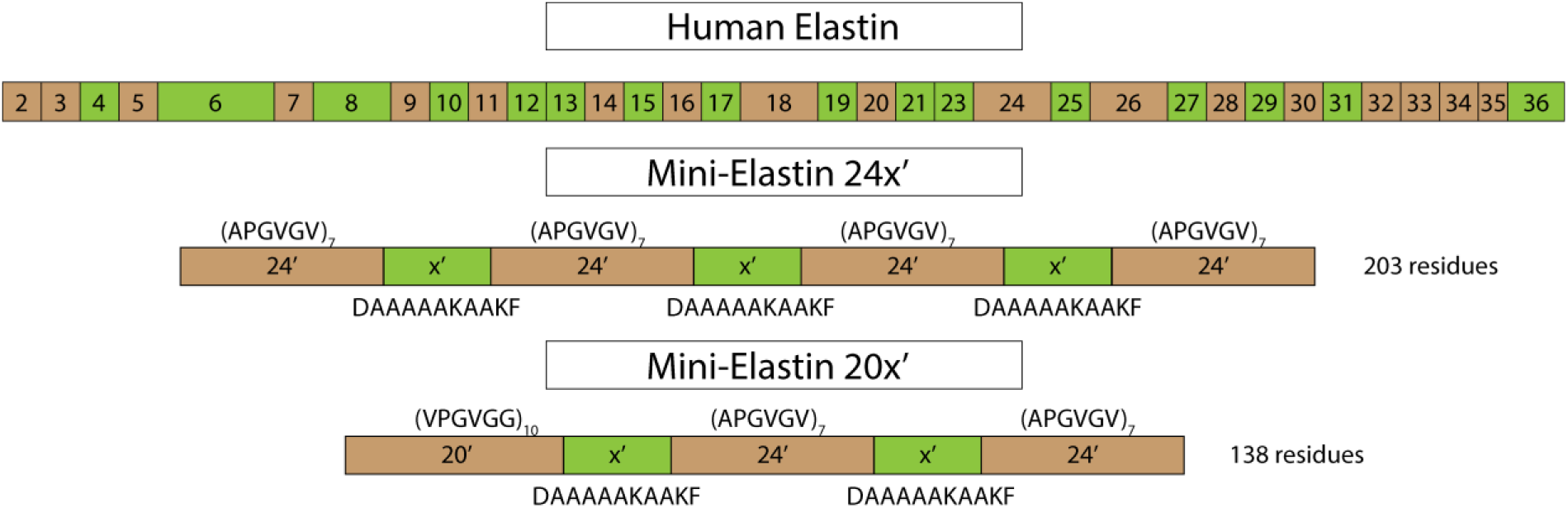
Domain structure of human elastin and the domain structure and sequences of minielastin proteins **24x′** and **20x′**. Hydrophobic domains are brown and crosslinking domains are green.

We then examined the natural, purified elastomer using solid state ^13^C NMR without magic angle spinning. Chain ordering was determined from the residual shielding anisotropy and dynamical timescales from R_1_ and R_2_ of the backbone carbonyl resonances. We find a similarly high degree of disorder in the natural protein and, importantly, mechanical stretching does not detectably decrease this disorder. As the polymer disorder we observe in both relaxed and stretched elastin is essentially identical, we infer that elastin’s recoil does not arise from configurational entropy loss at the residue level.

## THEORY

Dynamic analyses using R_1_, R_2_ and NOE data are less well developed for IDPs than for folded proteins. In both cases, the key function in the analysis is J(ω), the Fourier transform of the correlation function c(t) for the dynamics that contribute to spin relaxation. R_1_, R_2_ and NOE are related to J(ω) by three standard equations (SI eqs. 1a-c) (14, 16, 17). In the widely used Lipari-Szabo model-free approach (LS), c(t) is factored into independent dynamical modes, each of which is parameterized by an effective correlation time and a corresponding order parameter that are related to the timescale and amplitude of each dynamical mode, respectively (18, 19). LS has been used to analyze NMR relaxation in folded proteins and recently to analyze the 30-residue disordered terminus of an otherwise folded protein (14, 15). However, the general application of LS to IDPs has been questioned (20) and an alternative procedure, spectral density mapping (SDM), has also been used (21). In this method, the correlation function is not parameterized. Instead, the spectral density at five frequencies (0, ω_N_, ω_H_-ω_N_, and ω_H_+ω_N_) is determined from R_1_, R_2_ and NOE at two or more magnetic field strengths (20, 21). Insofar as SI eqs. 1a-c are valid, SDM is rigorous. However, SDM does not directly relate to molecular properties such as the timescales and amplitudes of dynamical modes that are discussed next in the context of parameterized spectral densities. A useful test is to compare the parameterized spectral density with the spectral density map.

An adjunct to SDM that potentially provides greater physical insight is the general form of the correlation function, c(t), for a dynamical process like diffusive or jump-like dynamics (22, 23):

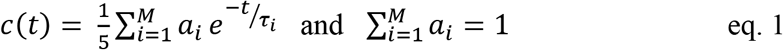

The spectral density is then

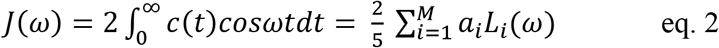

and 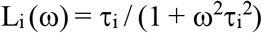 is the usual Lorentzian function with correlation time τ_i_. By constraining the correlation times to a range from 21 ps to 21 ns and separated by a factor of 4, Khan and co-workers have limited the general correlation function to 6 terms with 5 adjustable coefficients, ai (15). A method for detecting the predominant correlation times in the spectral density has also been recently described (24). Here, we simply limit the number of terms in eq. 1 so that the minimum number of parameters required to fit the data within experimental errors is not exceeded. In this way, the timescales of the dynamical modes present in the system under study can be identified. Note that the coefficients specify the contribution of each dynamical mode to the total spectral density but not a physical property like the amplitude of a motion. To better understand the coefficients, a master equation for a specific dynamical model can be used (22). However, for even simple models, the number of terms in the spectral density, eq. 2, typically exceeds what is experimentally accessible. Ways to reduce this in the context of structured proteins have been discussed in detail(18, 19, 25) and those potentially relevant to IDPs are summarized next.

For IDPs in solution, the slowest motion is diffusional reorientation of the aggregate protein. Insofar as the correlation times for overall reorientation are greater by a factor of 10 or more than those for internal dynamics, c(t) can be approximated as a product (19, 24, 26):

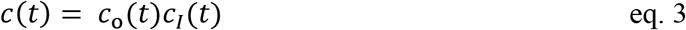

where c_o_(t) and c_I_(t) are, respectively, the correlation functions for overall and internal motions. For structured proteins that are not spherical, c_o_(t) is well approximated by a correlation function with two terms (19):

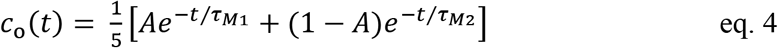

With A = 1, eq. 4 reduces to c_o_(t) for a spherical protein. Unlike structured proteins, IDPs have a flexible backbone and, in turn, a distribution of hydrodynamic radii. The distribution of correlation times is not easily obtained from NMR relaxation of backbone atoms which is dependent on both overall reorientation and large amplitude internal motions. Norton and coworkers circumvented this problem by assuming that the hydrodynamic radius, r_H_, in the Stokes-Einstein equations for translational diffusion, D_t_ = k_B_T/6πηr_H_, and rotational diffusion, 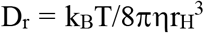, are the same (27). Importantly, Dt and thus the average hydrodynamic radius can be independently determined by PFG NMR or ultracentrifugation (6, 28). For IDPs with molecular weights comparable to **24x′**, the distribution of hydrodynamic radii is approximately gaussian with an average hydrodynamic radius, <r_H_>, that is found to be 4 to 7-fold greater than the gaussian width, σ so that negative hydrodynamic radii are excluded (29, 30). With this distribution, <r_H_^n^> and <r_H_>^n^ are equivalent within a few percent for n = −1, 1 and 3, and are used interchangeably herein. Thus, we calculate the average hydrodynamic radius from the translational diffusion constant:

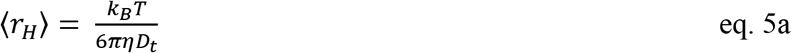

and, in turn, the average rotational correlation time is:

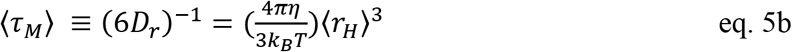

Averaging eq. 4 with a discreet gaussian distribution and A = 1, we obtain:

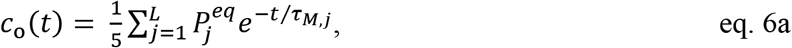

with:

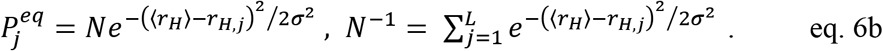

To account for internal dynamics with more than a single exponential correlation function, we use the 4-parameter cI(t) from the extended LS method (26):

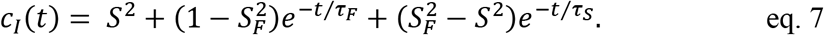

S_i_ are the generalized order parameters and τ_i_ are effective correlation times for fast (F) and slow (S) internal motions, respectively. The three order parameters are constrained by the relation S^2^= S_F_^2^S_S_^2^. In the limit of axially symmetric motion S = <P_2_(cosθ)>, which is the same as the order parameter S used in solid state NMR (19). When using eq. 7, it is assumed that the fast and slow modes can be approximated as a single exponential correlation functions and that τ_F_ and τ_S_ differ by an order of magnitude or more (26). With these conditions, the order parameters (S_i_^2^), are equilibrium properties of the dynamics.

The total correlation function, eq. 3, is the product of eqs. 6a and 7:

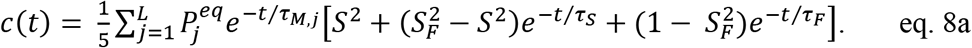

Eq. 8a has 3L terms and this simplifies to L+2 terms when overall reorientation is slow compared to internal dynamics, τ_M,j_ ≫ τ_S_ and τ_F_. After Fourier transformation, the result is:

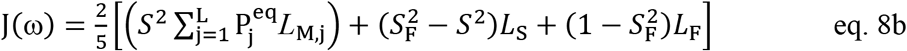

In the limit that τ_M,j_ are large compared to the reciprocal of the lowest NMR frequency (0.3 ns for the ^15^N angular frequency at the lowest field in this study), eq. 8b reduces to three terms:

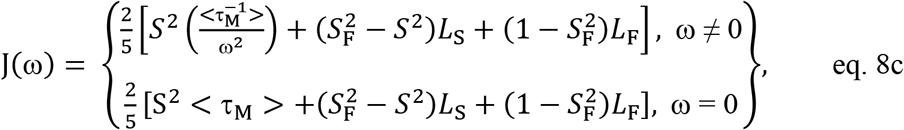

with:

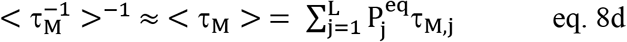

In this limit, eq. 1 truncated to three terms and eq. 8c are formally equivalent with parameters related by,

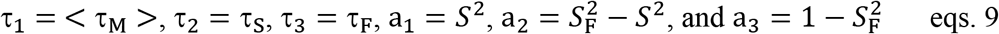

The above results can also be used to analyze relaxation in fibrous elastin, a material that is extensively cross-linked and overall reorientation is quenched. In the limit of τ_*M,j*_ → *∞*, eq. 8b reduces to,

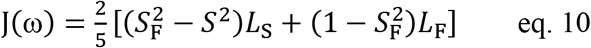

## RESULTS

### Backbone amide solvent exchange

To focus on motional dynamics, we first show that proton exchange at solvent exposed amides, known to affect NMR relaxation (31) is negligible in these experiments. Also, CEST experiments show the absence of other slow exchange processes. Proton exchange rates were obtained using the CLEANEX-PM method (32) (Figure 1). Experimental studies of an IDP and theoretical analysis by Baum and co-workers indicate that amide proton exchange rates of 10 s^−1^ to 30 s^−1^ increase amide proton R_2_ rates by 20–30%. In this study, relaxation data was obtained at pH 6, where exchange rates, Figure 1, are reduced to less than 2 s^−1^ and thus make negligible contributions to R_2_ (31).

**Figure 1.**
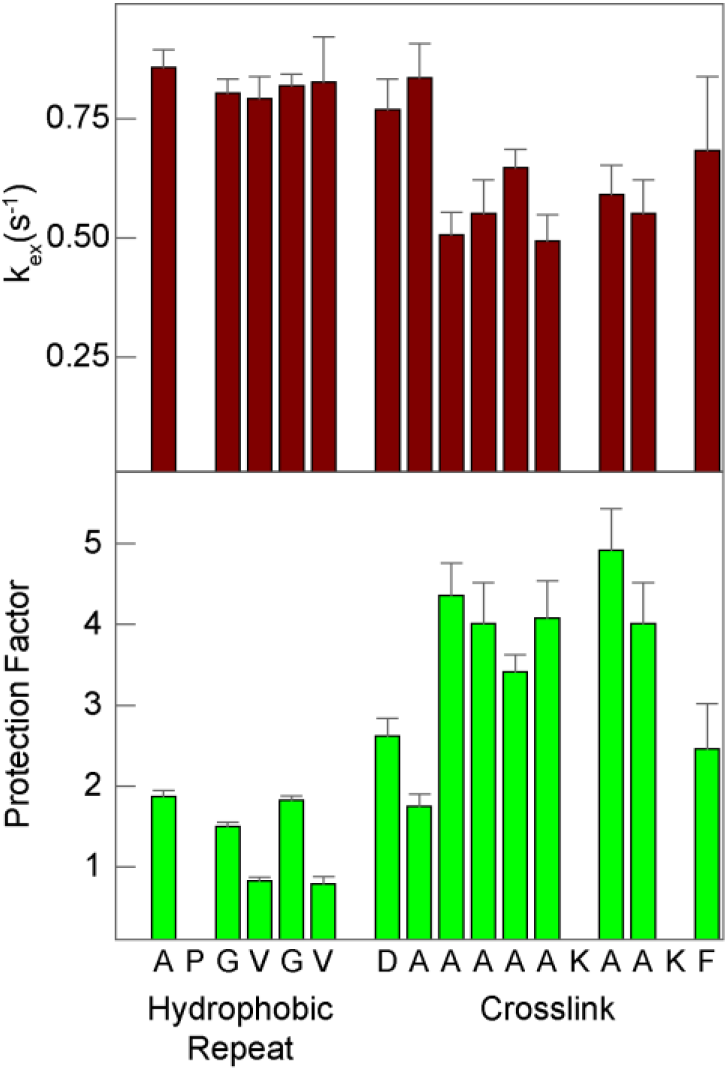
Amide proton exchange rates (top) in the hydrophobic repeat **24′**= (APGVGV)_7_, and cross-link, **x′**= (DA_5_KA_2_KF), modules at pH 6 in **24x′**. Protection factors (bottom) are calculated from the exchange rates as described by Englander and co-workers(33).

Protection factors calculated from the exchange rates (33, 34) are less than two for residues in hydrophobic modules and, systematically, higher than three for residues in the central portion of the cross-link module. Protection factors less than five indicate an absence of secondary structure (35). Thus, these amide protein exchange rates reflect the absence of secondary structure in the hydrophobic modules and weak ordering in the central residues of the cross-link modules confirming our previous conclusion formed on the basis of backbone chemical shifts (6).

*Backbone ^15^N and ^13^C relaxation parameters* obtained from **24x′**, a 203 residue minielastin, are shown graphically in Figure 2. ^15^N data was obtained at three NMR frequencies (500, 600 and 800 MHz) and ^13^C data was obtained at two frequencies (500 and 700 MHz). The complete data sets are listed in SI Tables 1a, c. ^15^N relaxation data from a 138 residue minielastin, **20x’**, closely parallels the ^15^N data from **24x’** and is listed in SI table 1b. There are three key features of the data. (i) The faster R_1_ and R_2_ relaxation rates and larger ^15^N NOEs observed in the cross-link modules indicate slower and/or more restricted motion than in the hydrophobic modules. This is consistent with our earlier chemical shift-based prediction of partial helical conformation in the cross-link modules (6). (ii) The small R_2_ / R_1_ ratios (~2 to ~3) in both hydrophobic repeat and cross-link residues indicate that fast backbone motions with correlation times less than a few ns affect spin relaxation significantly more than overall reorientation of the protein which has a substantially longer correlation time (τ_M_ > 10 ns) (14). (iii) Observed NOEs are frequency dependent and approach the slow-motion limit (NOE ~ 0.85) at 800 MHz. This indicates the presence of motions with correlation time(s) shorter than that for overall reorientation and longer than that for fast backbone motions (26).

**Figure 2.**
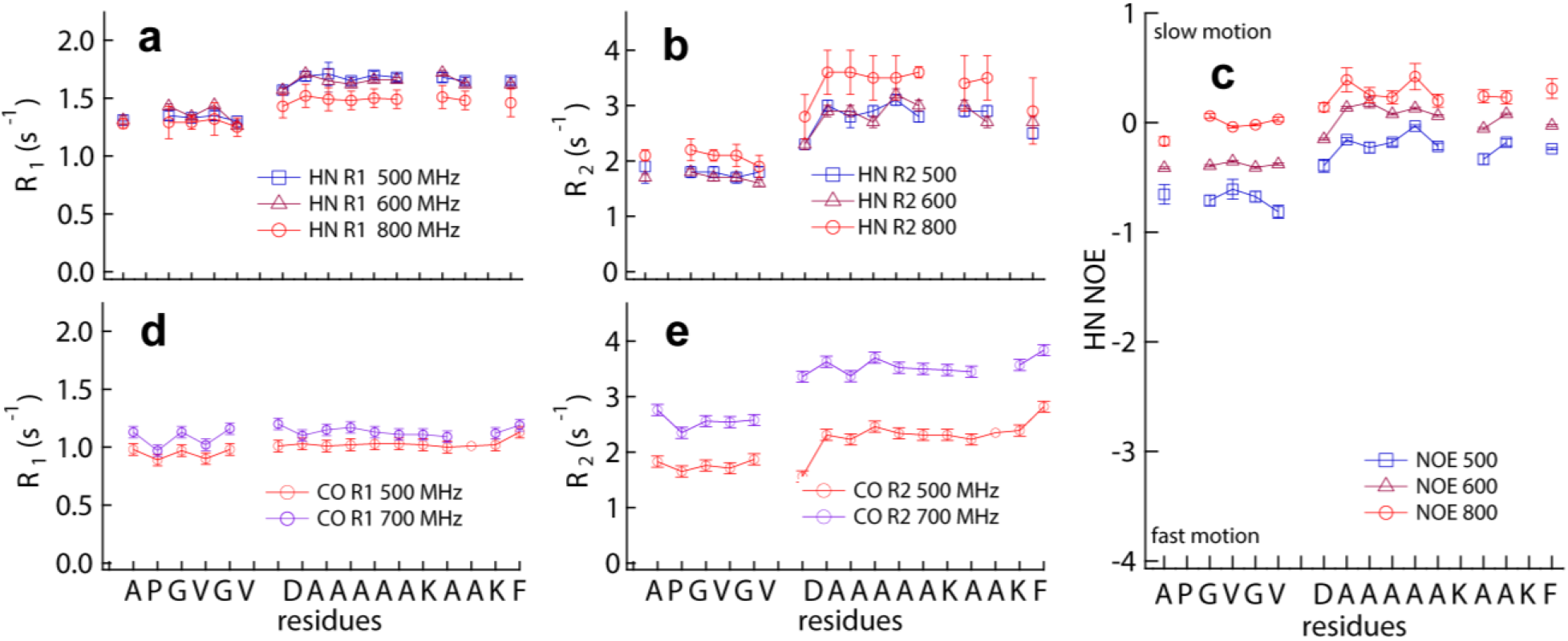
NMR relaxation data for **24x’** at the indicated NMR frequencies. (a-c) ^15^N amide R_1_, R_2_ and NOE. (d,e) ^13^C carbonyl R_1_ and R_2_.

*Fits of the ^15^N relaxation data* using eq. 1 truncated at 3 terms and equation 8a are summarized in Figure 3. While R_1_ and R_2_ values could be fit with two-terms in eq. 1, this 3-parameter correlation function predicted NOEs that were less than the experimental values. This was resolved by adding a third term with an intermediate correlation time and an additional coefficient (26). χ^2^ surfaces for this 5-parameter fit with eq. 1 are shown for a representative residue, A1, in the hydrophobic repeat, Figure 3a-c. Residue specific parameters from this fit are shown in Figure 3d-g (red marks). Standard errors are typically ± 15% except for the slowest motion that is fit with τ_1_ values from 4 ns to more than 50 ns and a1 from 0 to 0.15. However, within this large range, the best fit values of τ_2_, τ_3_, a2 and a3 are essentially unchanged (Figure 3b,c). Importantly, there is a time scale separation between τ_2_ and τ_3_: in all cases τ_2_ > 10τ_3_

**Figure 3.**
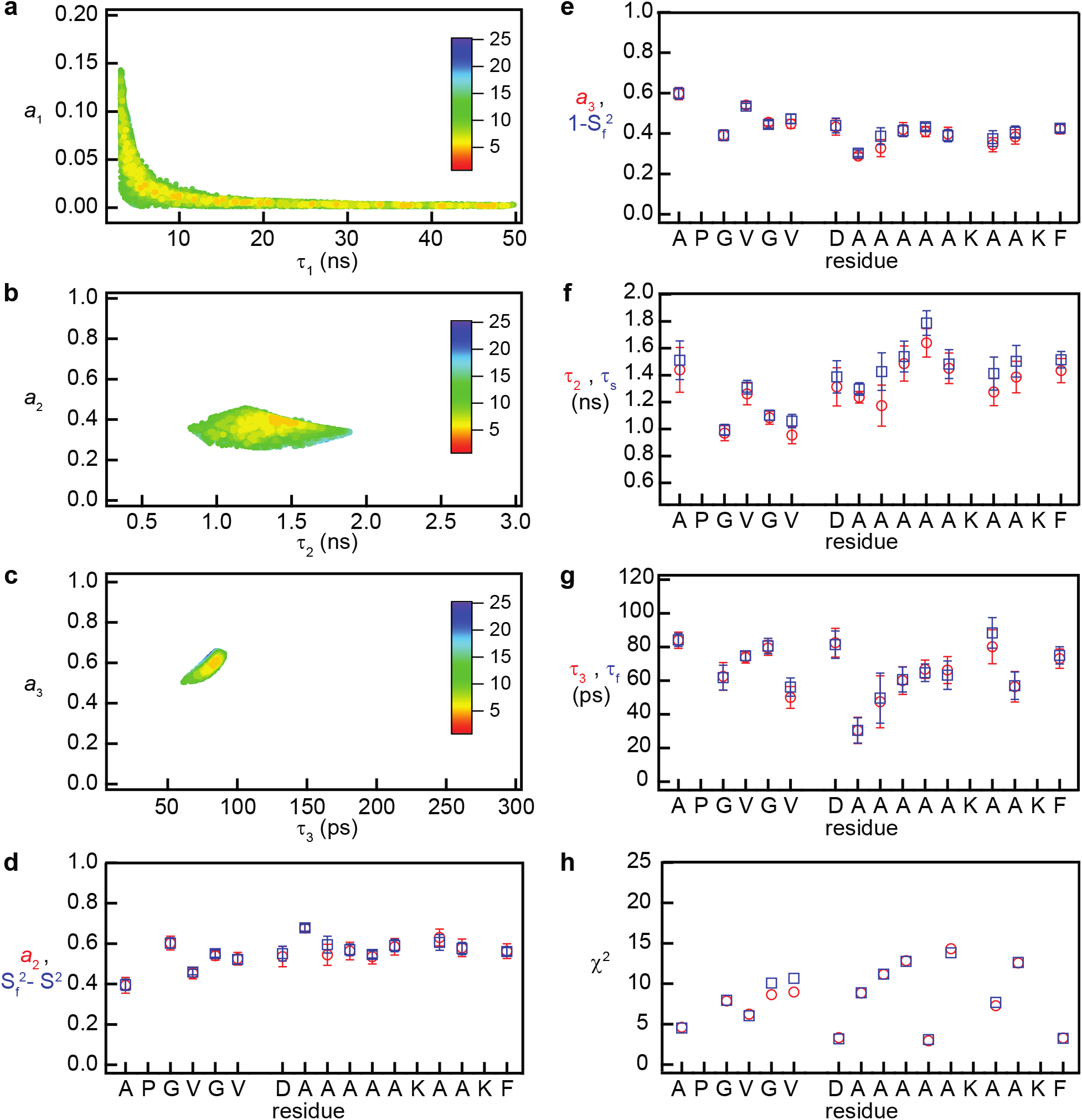
(**a-c)**χ^2^ surfaces for the fit of the eq. 1 spectral density to the relaxation data from the first residue of the hydrophobic repeat (A1). **(d-g, red)** Fit parameters obtained using the three term Lorentzian (5 parameters), eq. 1. **(d-g, blue)** Fit obtained using eq. 8 (4 parameters). (**h)** Per residue χ^2^ of the fits using eq. 1 (red) and eq. 8 (blue).

To determine the slowest motion, we have used the hydrodynamic radii previously determined by PFG NMR for **20x′** and **24x′**, 29.4 Å and 35 Å, respectively (6). Other possible slow motions, amide proton exchange and slow conformational change, are shown above to have negligible effect on R_2_ relaxation rates and we assume that the slowest motion in these soluble minielastins is global reorientation of the disordered protein. Using the Stokes-Einstein relation for rotational diffusion, the average rotational correlation times are 36 ns for **24x′** and 22 ns for **20x′**. Combined with the time scale separation between τ_1_ and τ_2_, we see that <τ_M_> > 10τ_2_ > 10τ_3_ and the modified LS spectral density is fit to the **24x′** relaxation data with <τ_M_> = 36 ns. Fits of the four adjustable parameters to eq. 8b, Figures 3d-g (blue marks) and SI Table 3, are insensitive to the width of the distribution of hydrodynamic radii and the spectral density 8b reduces to eq. 8c. Since the value of <τ_M_> determined from the hydrodynamic radius is within the large range that τ_1_ is constrained by the relaxation data alone, τ_S_ and τ_F_ are essentially equivalent to τ_2_ and τ_3_ and the minimum χ^2^ values, Figure 3h, for the 4-paramater fit are, in most cases, the same as for the 5-parameter fits. Values of the correlation times for slow internal motions (chain dynamic) vary from 1.0 ± 0.1 ns to1.5 ± 0.2 ns for residues in the hydrophobic repeat and 1.3 ± 0.1 ns to 1.8 ± 0.1 ns in the cross-link modules. Correlation times for the fast chain motions vary from 56 ± 5 ps to 84 ± 4 ps in the hydrophobic repeat and from 30 ± 7 ps to 88 ± 7 ps in the cross-link modules with the longer values grouped in the in the center of the module. Because of the time scale separation between parameters, order parameters 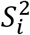 indicating the amplitudes of the backbone motions have been determined in addition to the correlation times. A striking result of this analysis is the nearly complete overall dynamic disorder, 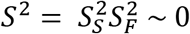.

To test these conclusions, we performed the same analysis of ^15^N relaxation data from **20x′,** a smaller minielastin (SI Table 1b), and to ^13^C relaxation data from the backbone carbonyl atoms in **24x′**(Figure 2d,e and SI Table 1c). Since **20x′**(138 residues) is shorter than **24x′**(203 residues), the correlation time for global reorientation is closer to the backbone correlation times and this experiment tests the assumption that global reorientation and backbone dynamics in an IDP can be treated as independent dynamical modes due to their different time-scales. The modular structure of **20x′**-20′-x′-24′-x′-24′(20′= (VPGVGG)_5_) -has the same cross-linker flanked by the same hydrophobic modules at the c-terminus and the chemical shifts of residues in the 24’ and x’ modules are equivalent to those in **24x′**. Fits of the ^15^N relaxation using the LS spectral density and the smaller rotational correlation time (<τ_M_> = 24 ns) yields backbone order parameters, *S*^2^ and *S*_F_^2^, and correlation times, τ_S_ and τ_F_, that are essentially equivalent to those determined for **24x′**. We conclude that the assumption of independent global reorientation and internal dynamics is a good approximation for minielastins with molecular weights greater than 13 kD. The ^13^C experiments examine backbone motions at atomic locations between the amide groups and sample the spectral density at different frequencies. From the analysis of the ^13^C data, SI Table 3c, we see that the correlation times and order parameters are in close agreement with those determined using ^15^N NMR, SI Table 3a. Thus, the high backbone disorder found at the backbone amide sites is also observed at the intervening carbonyl sites.

The agreement between the LS spectral density and the spectral density map, Figure 4, is excellent in frequency ranges where the two methods overlap and both analyses show that the dynamics of residues in the hydrophobic modules and the cross-link modules are different. Contributions to the parameterized spectral density from the three dynamical modes are shown in Figure 4b. At frequencies below 0.5×10^9^ s^−1^, the contribution from slow backbone motions is greater than the contribution from much slower global reorientation due to the high amplitude of the backbone motions (*S*^2^ ~ 0). At frequencies above 6×10^9^ s^−1^, only motions from fast backbone dynamics contribute to the spectral density.

**Figure 4.**
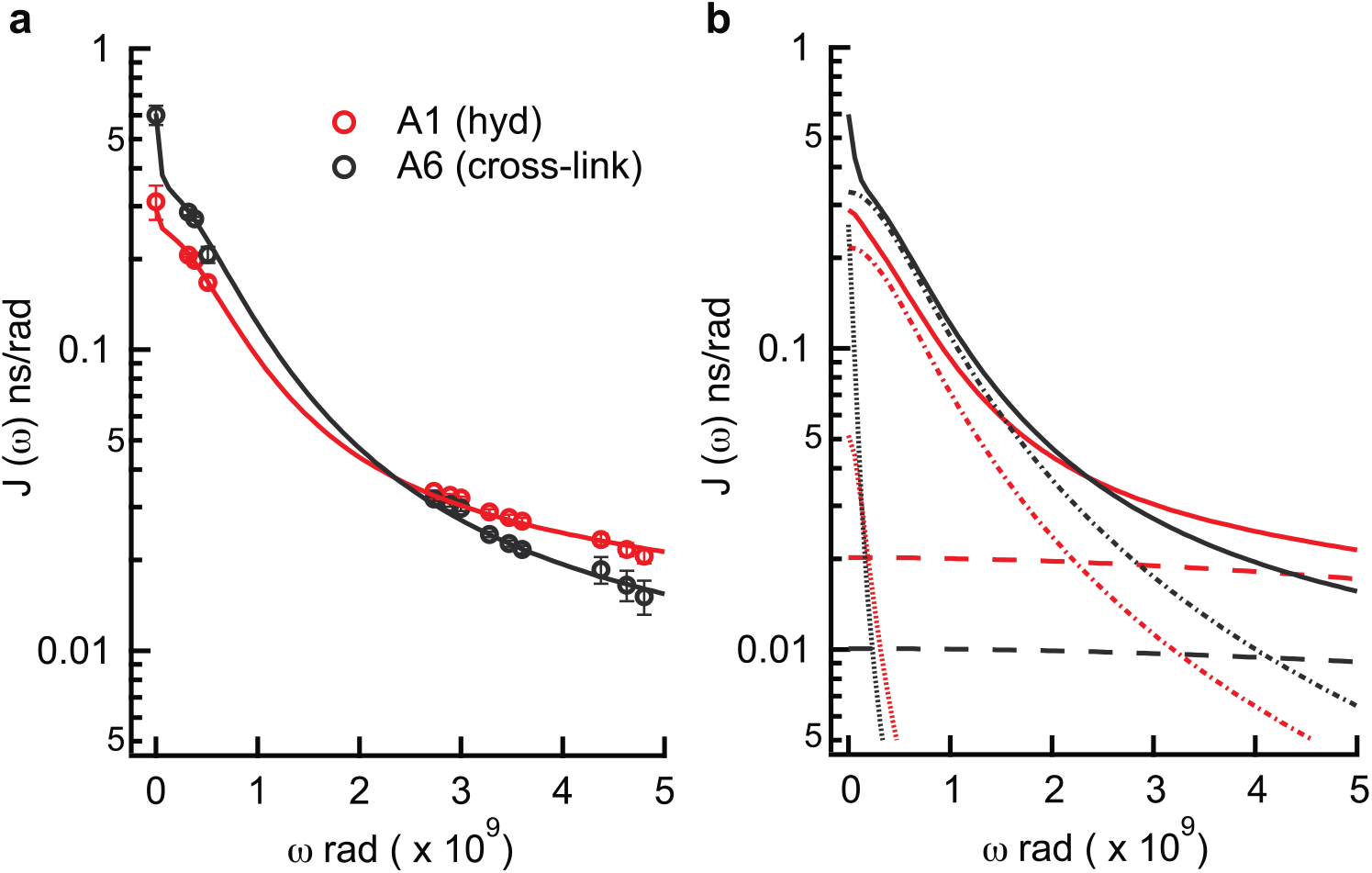
Logarithmic graphs of the spectral density versus NMR frequency. Red and black indicate A1 in the hydrophobic repeat and A6 in the center of the cross-link module, respectively. **(a)** The total spectral density obtained from eq. 8 with Figure 3 parameters (continuous curve) and SDM (filled circles). **(b)** Contributions to the total spectral density (continuous curves) from the terms with slow (∙∙∙∙∙), intermediate (∙ − ∙ −) and fast (− − −) correlation times.

### Dynamic analysis of purified bovine elastin

To compare dynamics in fibrous elastin with soluble **24x′**, we have studied the backbone carbonyl atoms of purified elastin fibers with ^13^C NMR. Carbonyl isotropic shifts are dispersed over a small range that is well separated from other carbon shifts and 98% of carbonyl groups in elastin are from the protein backbone. Since global reorientation of the highly cross-linked protein does not occur, the overall order parameter, *S*, for backbone motions can be estimated from the line width of spectra obtained without magic angle spinning (MAS). Linewidths in the spectra of static samples (Figures 5b-c) are 9-10 ppm and similar to those previously observed with MAS (9) and small compared to static shielding anisotropy of carbonyl groups, Δσ_stat_ ~ 116 ppm, indicating that the carbonyl groups in fibrous elastin are highly disordered. The small residual anisotropy and, in turn, the backbone order parameter were estimated from the spectrum line width by accounting for the contributions to the line width from the experimentally determined spin relaxation rate R_2_/π = 640 Hz (5 ppm) and the dispersion of isotropic chemical shifts. A simulated ^13^C spectrum of tropoelastin carbonyl isotropic shifts was convoluted with the experimentally determined R_2_ and then fit to a Lorentzian (Figure 5b,c). Isotropic shifts were simulated for the bovine tropoelastin sequence (36) using the chemical shift protocol that predicted observed ^13^C′ shifts in **24x′**(6, 37). The residual shielding anisotropy, Δσ_res_, is then the difference between the linewidths of Lorentzian fits of the observed and the calculated spectra indicating a residual anisotropy in the range of 1 - 4 ppm. In turn, the estimated value of *S* for chain dynamics in purified elastin, Δσ_res_/Δσstat, is in the range of 0.01 - 0.03 and *S*^2^ ~ .001 which is well within the range of *S*^2^ determined for the minielastins using ^15^N or ^13^C NMR relaxation. At the resolution of this experiment, we find no evidence for either stretch induced ordering of the elastin backbone or greater ordering of the protein in cross-linked elastin fibers compared to soluble minielastins.

**Figure 5.**
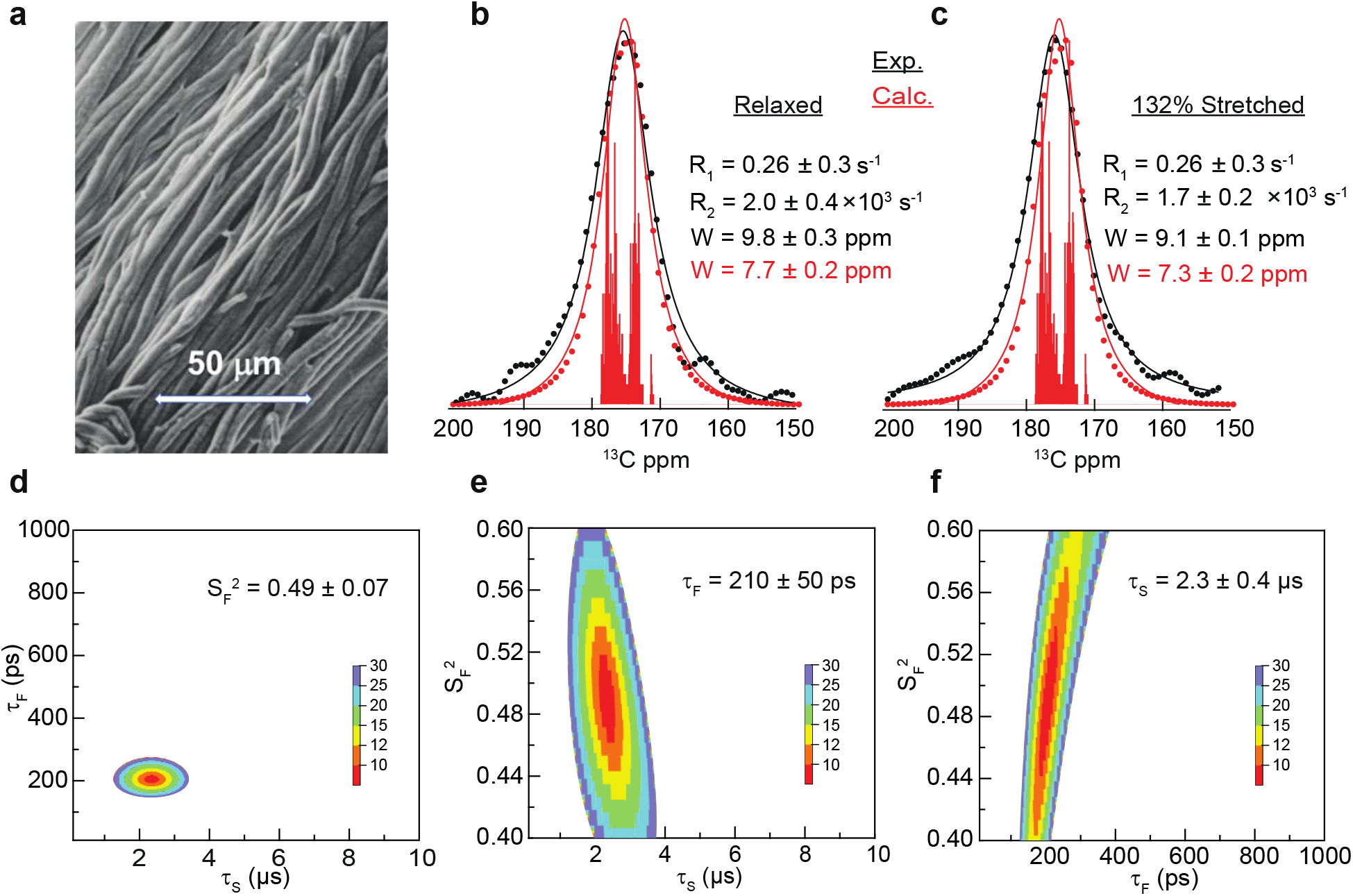
(a) SEM of purified bovine elastin fibers. (b, c) 500 MHz ^13^C spectra of (•) fibers relaxed and 132 % stretched. Vertical lines 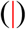 are the “stick” spectrum indicating the simulated chemical shifts of the bovine elastin sequence. Calculated spectra 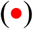 obtained by convoluting the stick spectrum with the indicated R_2_ values. Continuous curves are Lorentzian fits of the experimental (—) and calculated spectra 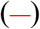 with the indicated linewidths, W. (d-f) χ^2^ surfaces of eq. 10 fit to the relaxation data (R_1_ and R_2_ at 500 and 700 MHz).

Combined with the order parameter determined from the residual shielding anisotropy of the backbone carbonyls, the timescales of backbone motions in natural elastin were determined from the 500 and 700 MHz R_1_ and R_2_ data, Figure 5 and SI Table 4. Compared to ^13^C relaxation rates of **24x′** in solution, R_1_ is four-fold lower and R_2_ is three orders of magnitude greater. Since global reorientation makes only a small contribution to relaxation in **24x′**(S^2^ ~ 0), these large differences in the relaxation rates are not due to the loss of global reorientation. Moreover, the observed R_1_ and R_2_ values are not consistent with a single correlation time for backbone dynamics and we have fit the relaxation data to equation 10, the modified Lipari-Szabo spectral density that includes backbone motion with two correlation times and no global reorientation. Since sample stretch did not affect R_1_ or R_2_ within experimental error, their averages were used and *S*^2^ was set at the value estimated from the residual anisotropy. The χ^2^ surfaces, Figure 5d-f, show that the correlation times τ_S_ and τ_F_ but not S_F_^2^ are well constrained by the available data. Limiting the fit range to 0.4 < S_F_^2^ < 0.6, we find that τ_F_ = 210 ± 50 ps and τ_S_ = 2.3 ± 0.4 μs. Compared to the soluble minielastins, elastin has no significant increase in backbone ordering, a large increase (by three orders of magnitude) in the slow correlation time for backbone motions and a small increase (four-fold) of the correlation time for the fast backbone motions. We also find no experimentally significant increase in backbone ordering when the elastin samples were stretched.

## DISCUSSION

We have used NMR methods to determine the timescales and the amplitudes of dynamics in two soluble minielastins and in purified elastin fibers. For the minielastins, ^13^C and ^15^N relaxation data were combined with PFG data and analyzed using three methods: spectral density mapping (21), a general spectral density function (22), and a modified Lipari-Szabo spectral density (18). The latter accounts for the distribution of hydrodynamic radii found in IDPs (29, 30) and protein backbone motions with well separated correlation times, τ_S_ and τ_F_. If the distribution of hydrodynamic radii is symmetric and the correlation time <τ_M_> for the average hydrodynamic radius is large, the modified Lipari-Szabo spectral density is mathematically equivalent to the general spectral density truncated at three Lorentzian terms. Analysis of the relaxation data obtained at three fields was found to be consistent with a general spectral density that has at least three Lorentzian terms and thus, the modified Lipari-Szabo spectral density as well. Fits of a two-term spectral density to the R_1_ and R_2_ data predicted incorrect NOEs. With three-terms, the correlation times τ_2_ = τ_S_ and τ_3_ = τ_F_ were well constrained by the relaxation data. At different residues, 1.0 < τ_S_ < 1.6 ns and 30 < τ_F_ < 84 ps. However, the longest correlation time, τ_1_, was not well-constrained by the NMR relaxation data alone (Figure 3a) indicating the need for additional information to determine the slowest motions. Effects on relaxation rates from slow conformational exchange and amide proton exchange rates were determined to be negligible indicating that the longest correlation time is due to overall reorientation of the protein, i.e., τ_1_ = <τ_M_>. Using the average hydrodynamic radius of 35Å for **24x′**(6) and the Stokes-Einstein relation, <τ_M_> was determined to be 36 ns. The separation of time scales, <τ_M_> > 20τ_S_ > 20τ_F_, indicates that a Lipari-Szabo type spectral density can be used and that the coefficients of the Lorentzian terms (ai in the general spectral density, equation 1) are related to generalized order parameters (*S*^2^and 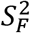 in equations 9) which are measures of the amplitudes of the “internal” chain dynamics. Although the number of different dynamical modes in an IDP is likely to be large, the number of terms in the spectral density was not extended beyond three given the available data. Previously, a three-term spectral density was used to analyze NMR relaxation data of an IDP (38). Herein, we have tested this simplification in three ways. First, the spectral density is in excellent agreement with the spectral density map over a wide range of frequencies indicating that it accounts for the available data, Figure 4. To test the key assumption that global reorientation of an IDP is independent of the large amplitude chain motions, NMR data from a smaller minielastin (138 compared to 203 residues) with a smaller <τ_M_> (24 ns compared to 36 ns) that is closer to the time-scale of the internal chain dynamics was also studied. The parameters (τ_F_, τ_S_, *S*^2^and 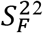) determined for the backbone motions in **20x′** from the relaxation data are the same as those for **24x′**. In other words, changing the length of the protein did not change the description of the chain motions obtained from our analysis. Finally, we used ^13^C NMR to study the backbone motions at the carbonyl groups in **24x′**. In this experiment, backbone dynamics were studied at positions close to the amide groups and the spectral density was sampled at additional frequencies. The correlation times and order parameters from our analysis of the ^13^C data, SI Table 3c, are in close agreement with those determined using ^15^N NMR, SI Table 3a.

Key results of our analysis is that the overall backbone order parameters, *S*^2^, at amide and carbonyl sites have been determined in addition to the correlation times; both of which can be compared to values found in folded proteins and other intrinsically disordered proteins. We find that the backbone correlation times in **20x′** and **24x′** are similar to those reported in both well-structured proteins (39) and in IDPs such as the disordered region of GCN4 (14), residues 146-199 of Engrailed 2 (15) and the disordered c-terminus of a Sendai virus protein (38). As expected, the overall order parameter for backbone motions in **20x′** and **24x′**, *S*^2^~0, is significantly smaller than found in well-structured proteins, *S*^2^~ 0.85 (17). However, the *S* ^2^ values in the minielastins studied here are also less than in the disordered region of GCN4 except for the most disordered residues at the N-terminus (14). Using equations 9 to interpret the coefficients of the three-term correlation function used in the analysis of the Sendai virus protein (38), we again find that this protein is also more ordered than **20x’ or 24x’** except for residues at the termini. We note that the disordered region of GCN4 folds upon DNA binding (40), whereas elastin retains high disorder after assembly into the mature elastic matrix. The absence of secondary structure in these minielastins that is indicated by NMR relaxation is in agreement with our previous results based on secondary chemical shifts and NOEs (6). IDPs with high proline content have a propensity for the formation of flexible structures with extended backbone conformations similar to polyproline II helices (28, 41, 42). For example, this is consistent with the observation of upfield C^α^ secondary shifts observed in Pdx1-C (43), an IDP with a proline content of 22%. However, upfield C^α^ secondary shifts were not observed in either **20x′** or **24x′** which have a smaller proline percentage in their sequences, 14% (6).

To compare dynamical amplitudes and timescales in soluble minielastins with those in mature cross-linked elastin fibers, we have refined an earlier study of disorder in bovine elastin (9) and determined the relevant correlation times in purified elastin fibers using natural abundance ^13^C NMR without magic angle spinning (MAS). While this experiment does not have residue specific resolution, it provides an overall picture of the timescale and amplitude of carbonyl group dynamics and refines an earlier estimate of backbone disorder obtained from ^13^C NMR with MAS (9). Due to extensive cross-linking, global reorientation of the protein is absent and the overall order parameter for chain dynamics is estimated directly from the contribution of residual ^13^C shielding anisotropy to the observed line widths. In both stretched and relaxed elastin, the residual anisotropy is small indicating that the average backbone order parameter in cross-linked elastin is essentially the same as found in soluble **20x′** and **24x′**. This result fully supports our analysis of the amplitude of backbone dynamics in the minielastins. Thus, elastin-like sequences are highly disordered both in solution and in the natural, cross-linked material (6, 9). Results from Reichfield and coworkers (44) indicate that this is also the case in a coacervated minielastin. However, the dramatically different relaxation times in elastin, Figure 5, indicate that backbone motions are significantly slowed in the cross-linked material: the slow correlation time, τ_S_, increases by a factor of 10^3^ while τ_F_ is much less affected and increases by a factor of 4. This suggests that the slow motion corresponds to chain dynamics on the length scale of the spacing between cross-links and the fast motion on a shorter and more local length scale that would be less affected by crosslinks. The spacing between cross-links in elastin is the length of the hydrophobic domains which varies from 11 to 55 residues (7). We conclude that structured regions in mature elastin are either absent or constitute a small part of the protein. Thus, a recent study of the naturally occurring cross-links in elastin which shows that all cross-link domains are connected in multiple ways (45) is in complete agreement with the dynamical properties of elastin and minielastins determined here.

## Conclusion

We find that the very high chain disorder observed in solution is retained in mature, cross-linked elastin. Moreover, no evidence for increased local ordering of the protein backbone induced by mechanical stretch is observed. While the exact quantitative correlation between backbone order parameters and configurational entropy remains a topic of discussion (46, 47), it is clear that order parameters as low as we observe in both minielastins and in natural elastin fibers in stretched and relaxed states are indicative of high flexibility and, consequently, high configurational entropy (48, 49). Previously, we reported a 10-fold increase of the ordering of water when fully hydrated elastin fibers are stretched by 50% (50). Together, these results support the hypothesis that stretch induced solvent ordering, i.e the hydrophobic effect, is a key player in the elastic recoil of elastin (11, 12).

## MATERIALS AND METHODS

### Protein expression

^15^N-labelled samples of minielastin constructs (Scheme 1) with sequence 24′-x′-24′-x′-24′-x′-24′ (**24x′**) and 20′-x′-24′-x′-24′(**20x′**), where **24′**= (APGVGV)_7_, **20′**= (VPGVGG)_5_ and **x′**= DA_5_KA_2_KF, were expressed and prepared as described previously (6). ^13^C,^15^N-labelled 24x’ was expressed using a 1/100 dilution of Bioexpress media (Cambridge Isotope Labs, Tewksbury, MA) in labelled M9 media (51). NMR samples were ~300 μM protein in pH 6, 50 mM phosphate buffer (90% H_2_O / 10% D_2_O).

*Purification of elastin fibers* from fresh bovine neck ligaments with a protocol (52) that yields pure elastin fibers with smooth surface and uniform diameter (Figure 5a). Briefly, samples of appropriate size for NMR were initially cut from the large ligament, washed with aqueous sodium chloride to remove water-soluble proteins and then with organic solvents to remove lipids. Other proteins were removed by treatment with cyanogen bromide (elastin has no methionine) followed by a wash with aqueous urea containing β-mercaptoethanol and final purification by limited trypsin digestion for 4 hours at 37 °C to remove micro-fibrillar components (in mature elastin, most trypsin cut sites are liminated due to cross-linking). Purified samples were stored at – 80 °C.

### NMR measurements of backbone amide exchange

Amide ^15^N-^1^H exchange rates in **24x′** were measured on a 700 MHz Varian Inova instrument using the CLEANEX-PM pulse sequence during which buildup of NMR signal, I(τ), as a function of the exchange time, τ, is (32):

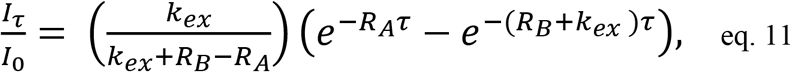

where I_0_ is the reference signal intensity obtained from the fast HSQC spectrum and kex is the amide proton exchange rate. RA and RB are NMR relaxation rates of the water and amide protons, respectively. Spectra were accumulated with 1500 and 64 complex points in the direct and indirect dimensions, respectively, 16 scans per t1 increment, a recycle delay of 1.5 s and 6 exchange times, 5, 50, 100, 200, 350 and 400 ms. Values of kex were extracted from the plots of the signal intensity ratio I(τ)/I_0_ versus τ and reported errors are errors of the fit.

*NMR relaxation measurements of minielastins* were collected at 298 K on Bruker Avance 500, 600, 700 and 800 MHz instruments equipped with cryoprobes. For ^15^N HN relaxation, 90° pulse widths on all four instruments varied from 7-10 μs (^1^H) and 25-40 μs (^15^N). Data were processed using NMRPipe (53) and TopSpin 3.5pI7 software. Relaxation rates were calculated using peak heights and steady-state [^1^H]-^15^N NOE values were calculated from the ratio of peak heights in NMR spectra acquired with and without proton saturation. The signal-to-noise ratio in each spectrum was used to estimate the experimental uncertainty.

^15^N R_1_, R_2_ and NOE at 600 MHz were measured by chemical exchange saturation transfer (CEST) (54) to probe for the presence of slow exchange processes, which are not present. ^15^N CEST spectra were recorded with a B1 field of 87.5 +/− 4 Hz calibrated as previously described (55, 56), with an ^1^H decoupling field strength of 3.5 kHz centered at 8.5 ppm during the mixing time (500 ms) and 70 B1 offsets from 102 to 136.5 ppm. A reference experiment with a null mixing time was also acquired. The spectral parameters were 512 and 64 complex points in the direct and indirect dimensions, respectively, 4 scans per t1 increment and a recycle delay of 1.5 s. Data were plotted as normalized peak intensity (to the reference intensity) vs. offset and errors in the data points were estimated from the deviation of peak intensities where peak attenuation was not occurring. Extraction of R_1_ and R_2_ from the data utilized an in-house python script. R_1_ spectra were recorded with ten delay times (10 to 1200 ms) and 2 s recycle delay and R_2_ spectra with eight delays (10 to 350 ms) and 1 s recycle delay. NOE spectra were recorded with a 10 s saturation period and 4 s recycle delay. Acquisition parameters were 1024 and 200 complex points in direct and indirect dimensions, respectively, and 20 scans per t_1_ increment.

^15^N R_1_, CPMG R_2_ and NOE at 500 and 800 MHz on **24x’** were determined using the pulse sequences of Farrow et al. (57). R1 spectra were recorded as pseudo-3D experiments with nine delay times (10 to 1200 ms) and 2 s recycle delay; R_2_ pseudo-3D spectra were acquired with eight CPMG delay times (17 to 340 ms) and a 1 s recycle delay. NOE spectra were recorded with a 5 s saturation period and no recycle delay.

^15^N R_1_, R_2_ and NOE at 500 and 700 MHz on **20x’** were determined using the pulse sequences of Farrow et al. (57). R_1_ spectra were recorded as pseudo-3D experiments with nine delay times (20 to 1200 ms) and 1.5 s recycle delay; R2 pseudo-3D spectra were acquired with eight CPMG delay times (17 to 340 ms) and a 1.5 s recycle delay.

HNCO-based ^13^CO R_1_ and R_2_ at 500 and 700 on **24x’** were determined using the pulse sequences of Chang and Tjandra (58). R1 spectra were recorded with nine delay times (10 to 1600 ms) and a 1.5 s recycle delay; R1rho spectra were acquired with a B1 field of 2500 Hz, seven tau values (10 to 240 ms), and a 1.5 s recycle delay.

*Fitting parameters (correlation times and order parameters) to the relaxation data.* R_1_, R_2_, and NOE were fit with SI eqs. 1a–c and spectral density functions eq. 1 or 8b for minielastin and eq. 10 for purified natural elastin fibers. To account for the contributions to relaxation from dipolar coupling to protons and chemical shielding anisotropy, we have used rHN = 1.02 Å, η = 0 and δ’_z_=−114.7 ppm for ^15^N and rHC′ = 1.69 Å (14), η = 0.81 and δ’_z_ = −77 ppm for ^13^C′ (59). Optimum parameter values were obtained by a Monte Carlo or a grid search for the minimum value of χ^2^ with χ^2^ defined (60, 61) as:

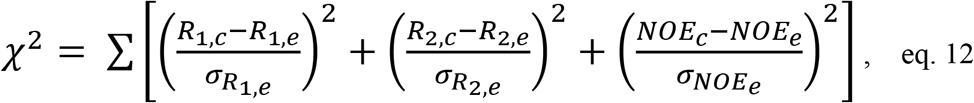

and subscripts “e” and “c” indicate experimental and calculated values, respectively. The sum is over NMR frequencies. Standard errors in the parameters were determined by Monte Carlo simulation as previously described (62–64). Five hundred normally distributed data sets were generated using the relaxation parameters back calculated from the optimum fit and the experimental uncertainties (SI Tables 1 and 4) as the gaussian means and standard deviations, respectively. A distribution for each fit parameter was then obtained by fitting each of the data sets in the same way as the experimental data. The fit parameters, shown in Figures 3 and 5 and listed in SI Tables 2, 3 and 5, are the optimum parameters obtained at minimum χ^2^ with the indicated error limit calculated as the standard deviation of each parameter distribution.

*Static ^13^C NMR spectra of stretched and relaxed Elastin* were obtained on a homebuilt 500 MHz spectrometer and a 700 MHz Varian Inova spectrometer. For the 500 MHz instrument, the ends of a dry elastin sample (~ 2 mm × 25 mm) were super-glued to ~2 cm lengths of 3.2 mm G10 rod. Following overnight equilibration in ^2^H_2_O, the fiber assembly was inserted into a 5 cm length of a 5 mm NMR tube open at both ends. At one end, the rod was sealed to the glass tube with glue. So that the sample could be stretched, the rod protruding at the other end was held in place and sealed with parafilm. This assembly was inserted into the horizontal coil of the homebuilt probe(65). R_1_ and R_2_ relaxation times were measured using inversion recovery and Hahn echo pulse sequences with relaxation delays of 4 s. An arrangement similar to the above was adapted for the unmodified cryoprobe of the 700 MHz instrument and a 5 mm sample tube with the closed end removed. G10 rods were cut to protrude from the ends of the NMR tube so that the elastin sample (2 mm × 29 mm) was centered in the sample coil. The lower rod was glued to the NMR tube and the upper rod was held in place with parafilm. R_1_ spectra were recorded at nine delays (0.25 to 12 s) and R_2_ spectra measured with a Hahn echo at 500 MHz and with CPMG echoes at 700 MHz were recorded at ten delays (0.2 to 1.2 ms) with 4 s recycle delays. The ^13^C NMR relaxation data of hydrated elastin was processed in a same manner as the **24x′** using χ^2^ minimization to obtain the best fit parameters, τ_F_, τ_S_ and S_F_^2^.

## ASSOCIATED CONTENT

### Supporting Information

All reference to the supporting information is included in the main text. The following files are available free of charge.

R_1_, R_2_ and NOE equations, relaxation data and fit parameters for **24x′,** CLEANEX data and relaxation data of purified elastin relaxed and stretched (PDF)

## Author Contributions

F.C.A.C., J.M.P., R.J.W., and R.L.K designed research; F.C.A.C., J.M.P., N.M.J., T.M.S., and J.M.A. performed research; F.C.A.C., R.J.W. and R.L.K wrote the manuscript.

## NOTES

The authors declare no competing financial interests.

## ACKNOWLEDGMENT

The authors gratefully acknowledge support from the NSF (DMR-1410678) to R.J.W and R.L.K. Program and infrastructure support was from the National Institutes of Health National Center for Research Resources to the City College of New York (5G12-MD007603-30). Prof. R. Koder is a member of the New York Structural Biology Center. Some of the work presented here was conducted at the Center on Macromolecular Dynamics by NMR Spectroscopy located at the New York Structural Biology Center, supported by a grant from the NIH National Institute of General Medical Sciences (GM118302). J.M.P. was the recipient of a fellowship award from the U.S. Department of Education Graduate Assistance in Areas of National Need (GAANN) Program in Biochemistry, Biophysics, and Biodesign at The City College of New York (PA200A150068).

## ABBREVIATIONS

CEST: chemical exchange saturation transfer
IDP: intrinsically disordered protein
NMR: nuclear magnetic resonance
NOE: nuclear Overhauser effect
PFG: pulsed field gradient
MAS: magic angle spinning
SDM: spectral density mapping

## Notes

### Competing Interest Statement

The authors have declared no competing interest.

### Summary of Updates

We have performed a significant number of additional experiments, including a full amide relaxation analysis of a differently sized but related minielastin protein and the relaxation analysis of backbone carbonyl relaxation in the original minielastin to allow direct comparison with the natural elastin fiber NMR data. We have further performed a significantly more detailed numerical analysis of our results.

